# Treadmill Stepping in Newborn Rats

**DOI:** 10.64898/2026.03.02.709074

**Authors:** Aimee L. Bozeman, R. Blaine Kempe, Nancy Devine, Tiffany S. Doherty, Dan Tappan, Misty M. Strain, Michele R. Brumley

## Abstract

The purpose of this study was to investigate the influence of treadmill belt speed on mechanically-induced (tail-pinch) and pharmacologically-induced (quipazine, a 5HT_2A_ agonist) stepping behavior in one-day old rats. On postnatal day 1 (P1), male rat pups were tested on one of four moving treadmill belt speeds. Stepping was induced using a tail-pinch and quipazine administration to examine real-time adaptations on belt speeds. For tail-pinch-induced stepping in the forelimbs and hindlimbs, there was a significant effect of time but not belt speed. Step cycle duration was significantly shorter for both forelimbs and hindlimbs on the fast belt speed compared to all other belt speeds. For the forelimbs, this effect was driven by changes in stance phase duration. Compared to control the speed, step area was significantly larger on medium and fast speeds for forelimbs and slow and fast speeds for hindlimbs. For quipazine-induced stepping, forelimbs and hindlimbs showed significantly more steps across slow, medium, and fast belt speeds compared to the control speed. The forelimbs showed significantly shorter step cycle durations on the fast belt speed compared to the control belt speed. Again, this difference was driven by changes in the stance phase. There were no significant differences in step cycle duration, stance and swing phase durations, or step area between speeds for the hindlimbs. Overall, we showed that both mechanical and pharmacological stimulation is reliable at inducing stepping on a moving treadmill belt in neonatal rats, and P1 rats show real-time adaptations in response to a moving treadmill belt.

## Introduction

Rodents are commonly used in studies investigating neurodevelopmental processes and neurobiological mechanisms of locomotion (i.e., Mistretta, Wood, English, & Alvarez, 2024; Vinay, Ben-Mabrouk, Brocard, Clarac, Jean-Xavier, Pearlstein, & Pflieger, 2005). In the rat, the neural mechanisms supporting locomotion develop well before quadrupedal locomotion is observed at approximately two weeks after birth (Altman & Sudarshan, 1975; Swann & Brumley, 2019). Previous research has shown that locomotor spinal networks develop prenatally, continue developing after birth in parallel with postural mechanisms, and are not fully developed until the third postnatal week (Gramsbergen, 1998; Swann & Brumley, 2019; Vinay, Brocard, Clarac, Norreel, Pearlstein, & Pflieger, 2002). For instance, *in vitro* studies with the isolated spinal cord of rats and mice have shown that thoracic and lumbar networks produce rhythmic hindlimb locomotor rhythms in the absence of movement (called “fictive locomotion”) before birth (Branchereau, Morin, Bonnot, Ballion, Chaprom, & Viala, 2000; Nishimaru & Kudo, 2000), and that locomotor-like stepping behavior can be induced in fetal rats (Brumley & Robinson, 2005). Importantly, the rodent model also continues to be the most prevalent system for studying spinal cord injury (Sobolev, Sysoev, Vyunova, & Musienko, 2025).

In behavioral and neurological studies, treadmills are often used for motor training and sensory stimulation to influence locomotor neurorehabilitation (Hagen & Fling, 2026; Hubli & Dietz, 2013). Sensory stimulation of the moving treadmill belt not only helps to initiate leg muscle activity necessary for locomotion; it also modulates stepping behavior over the training period. Classic experimental work with spinal-injured cats has shown that treadmill training can be used to ameliorate the behavioral and physiological effects of a spinal cord injury. The changes induced by treadmill training are seen in the quality of locomotion, inhibitory processes, and cellular properties (Edgerton et al., 1997; Tillakaratne et al. 2002). Early treadmill research with kittens has shown that developing cats are responsive to treadmill stimulation during the early postnatal period as well (Bradley & Smith, 1988), although they often require additional stimulation such as a sustained tail pinch to activate stepping. While many treadmill studies have been conducted using rodents, from investigating recovery following neural injury to therapeutic impacts on depression and obesity, no studies have evaluated if treadmill stimulation can be used to investigate neurodevelopment of locomotion during the neonatal period. Most research using treadmill training as a sensory stimulus has focused on restoration of motor function following a spinal cord injury or other neural insult. Often, these studies use rats that have sustained injuries in adulthood and then are trained on a treadmill to assess the efficacy of motor training paradigms for the recovery of locomotion following injury (Fouad et al., 2000; Multon et al., 2003; Cha et al., 2007). For example, one study found that rats with a complete spinal cord transection showed functional recovery of stepping in adulthood after being step-trained on a treadmill beginning at P31 (Tillakaratne et al., 2010).

It is well known that neonatal rats are responsive to environmental cues and many classes of sensory stimuli, including mechanical stimulation to evoke motor responses. These cues and stimuli can induce a variety of motor behaviors, activating the circuitry needed to produce and modulate these movements. For example, stimulation of the anogenital region via a vibrotactile device produces a reflexive leg-extension response, a coordinated motor behavior involving the bilateral extension of the hindlimbs, in postnatal day 1 (P1), P5, and P10 rat pups (Kauer, Allmond, Belnap, & Brumley, 2016). Following administration of a serotonin agonist, a tail pinch evokes stepping behavior in P1 rats, showing that intact, neonatal rats not only respond to sensory stimulation but also show mature stepping movements well before quadrupedal locomotion is observed (Swann, Kauer, Allmond, & Brumley, 2016). Additionally, a tail pinch induces short bursts of fictive locomotion in neonatal rats with a spinal cord transection (Lev-Tov, Etlin, & Blivis, 2010). Also, fetal studies using interlimb yoking procedures provide evidence that rats *in utero* are capable of motor learning before the nervous system is fully developed (Robinson, 2005; Robinson, Kleven, & Brumley, 2008). Moreover, it has been shown that the spinal cord mediates interlimb motor learning and memory effects in fetal rats (Robinson, 2015). Therefore, it seems plausible that rats shortly after birth might be able to respond to treadmill stimulation and modulate their motor behavior based on characteristics (i.e., speed) of the treadmill belt.

We started with the question of whether or not neonatal rats would exhibit stepping behavior in response to a moving treadmill belt (Brumley et al., 2015). However, we found that the moving treadmill belt alone was not sufficient for *inducing* stepping behavior. Here, in the present study, we asked if one-day old (P1) rat pups would *modulate* their stepping behavior in response to different treadmill belt speeds, if we first induced stepping via another mechanism (i.e., pharmacological or mechanical stimulation). Using neonatal rats, previous research from our lab has shown that quipazine, a 5-HT_2A_ agonist, evokes sustained interlimb stepping and locomotion (walking) behavior, and that a tail pinch induces an immediate, brief bout of alternating hindlimb stepping (Swann, Kauer, Allmond, & Brumley, 2017; Swann, Kempe, Van Orden, & Brumley, 2016). Thus by evoking stepping behavior using these methods, here we examined how speed of the treadmill belt speed would *modulate* ongoing stepping behavior in neonatal rats, by measuring step frequency, characteristics of the step cycle (swing and stance), and step area. If rats adapt their treadmill stepping to differences in belt speed shortly after birth, this would suggest that developing neural circuits for locomotion are already quite sophisticated in terms of sensorimotor regulation.

The purpose of this study was to investigate the influence of treadmill belt speed on tail-pinched induced, and then quipazine-induced, stepping behavior in one-day old rats. In this experiment, P1 rats were tested on a moving treadmill belt in one of four speed groups: control (non-moving), slow, medium, or fast. We hypothesized that rats in the non-moving belt speed group would show the lowest frequency of steps, followed by the slow and medium groups, with the fast group showing the highest frequency of steps. To accommodate changes in step frequency, we expected that parameters of stepping (limb kinematics) would also change. Because all four paws were placed on the treadmill, we examined changes in both the forelimbs and the hindlimbs. If neonatal rats show modulation of stepping behavior on a treadmill, that would suggest that this tool could be used for experimental investigation into neurobiological and developmental mechanisms of sensorimotor integration and recovery of function in immature rodents.

## Methods

### Subjects

Subjects were neonatal offspring of Sprague-Dawley rats housed in the Animal Care Facility at Idaho State University. Testing was conducted with 32 male P1 rat pups (∼24 h after vaginal delivery). Pups stayed with the dam in the home cage in the colony room on a 12:12 hour light:dark cycle until the time of testing. Ambient light, temperature, and humidity were controlled in the colony room. Before testing, each subject was examined for overall health by identifying a milk band on the belly (evidence of recent feeding), pinkish color, and healthy weight. No more than one pup per litter was assigned to each group in order to avoid litter effects. Care and use of all animals were in accordance with guidelines set by the Idaho State University Animal Care and Use Committee and the National Institute of Health guidelines (National Research Council, 2011).

### Experimental Design

#### Treadmill testing

Testing occurred inside an infant incubator with a controlled humidity of 40% and temperature of 35°C. Immediately prior to testing, rat pup subjects were manually voided using perineum stimulation, then weighed and placed in the incubator for 30-min of acclimation to testing conditions. Following acclimation, subjects were secured in the prone posture to a horizontal bar with a rubber surface, using a harness to gently hold the head and torso while the limbs hung down and were able to move freely (Brumley, Roberto, & Strain, 2012). Securing the animals at this age was necessary because newborn rats have relatively weak muscles and poor postural control. The bar was placed just above a miniature treadmill so that the hindlimbs and forelimbs of the subject made full plantar contact with the belt. Suspension from the bar also ensured that the subject stayed at the same height from, and at the center of, the treadmill without slipping or moving off. An illustration of a newborn rat subject on the bar, suspended above the treadmill, is provided in Figure 1. A small ruler was attached to the horizontal bar (but did not touch the subject) for calibration for later kinematic analysis.

**Figure 1.**
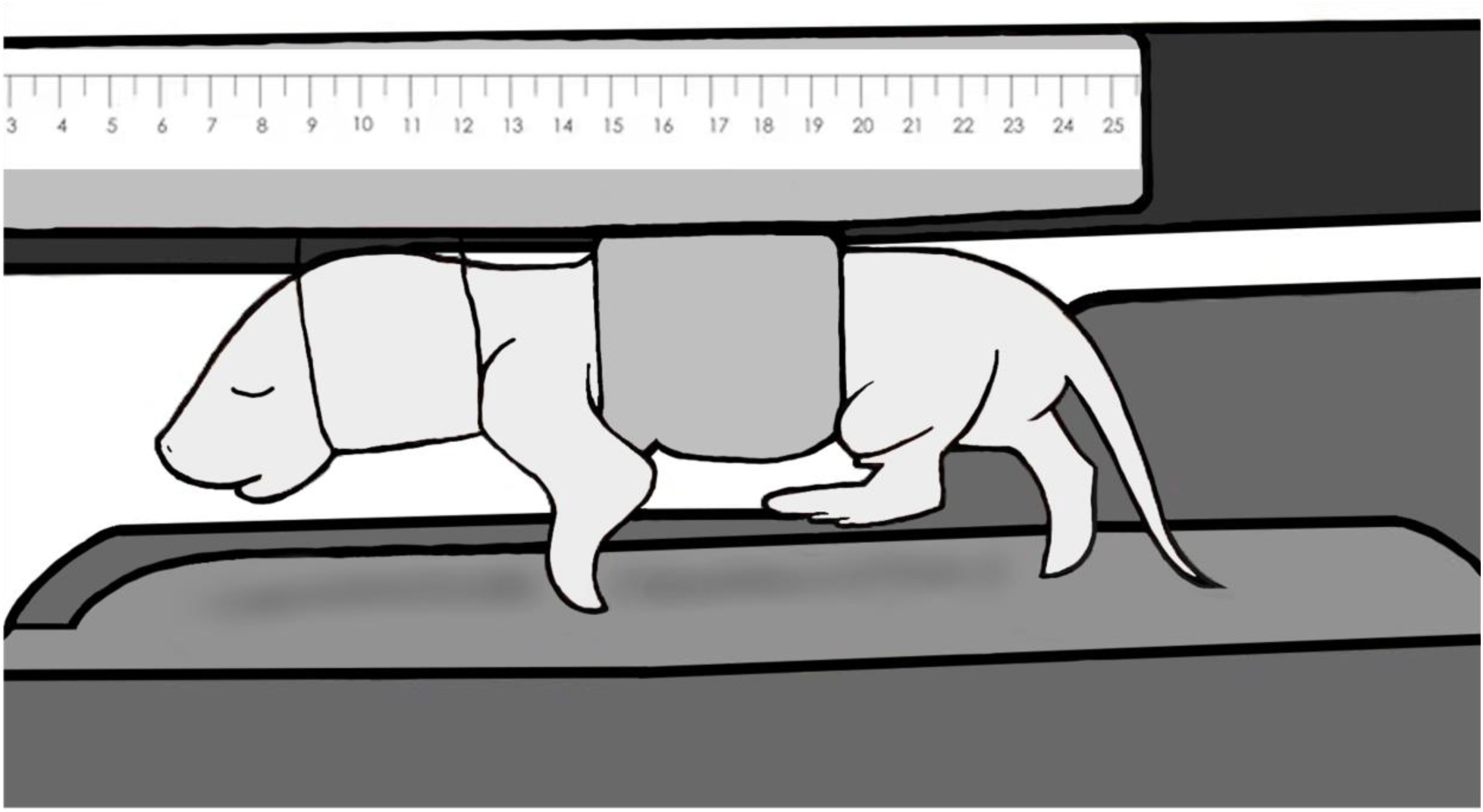
Illustration of treadmill set-up. The rat pup is secured to a support bar with medical tape around the abdomen and clear elastic around the head and neck. Limbs were able to move freely, and both fore- and hindpaws made full plantar contact with the treadmill belt. For step area analyses, a small ruler was secured to the support bar.

The dimensions of the treadmill belt were 9.5 cm in length and 4.5 cm in width. The length and width of the belt were large enough to prevent the limbs coming in contact with the edge of it. A soft, semi-elastic material (latex dental dam) covered the belt surface. The treadmill was connected to a computer via USB and was controlled by software designed by one of the authors (DT). Four belt speeds were used for the different conditions; non-moving (control), slow speed (1.72 cm/s), medium speed (2.46 cm/s), and fast speed (3.19 cm/s). The medium speed (2.46 cm/s) was calculated by measuring the average speed of a one-day-old rat’s alternating stepping limb movements following treatment with quipazine (Swann et al., 2016). The fast and slow speeds were calculated by increasing and decreasing the medium speed by 30%. Subjects were tested at only one belt speed.

#### Timeline of study

Each subject received tail-pinch (physical) stimulation and quipazine (pharmacological) stimulation, with one following the other by a 15-min inter-stimulation interval. Thus, we evoked stepping with tail pinch and then with quipazine and examined the effect of treadmill belt speed on stepping behavior following each form of stimulation.

After the subject was secured to the horizontal bar and placed onto the treadmill, the session started with a 1-min baseline. The purpose of the baseline was to acclimate the subject to the bar and treadmill. Following baseline, a tail pinch was administered. The effects of tail pinch in newborn rats subsides very quickly, typically with a robust behavioral response occurring within 15 sec, and a marked decrease in activity by 2 min after stimulation (Swann et al., 2017). Five min after tail pinch, the subject was raised from the treadmill, while still secured to the bar, for 5 min of rest. The subject was then lowered back onto the treadmill for a second baseline period, followed by administration of quipazine and a 30-min test period. Previous research shows that following quipazine administration alternating stepping in P1 rats significantly increases within 5 min and continues for at least 30-min post-injection (Swann et al., 2016). The entire session was recorded onto DVD. Other than for the control condition, the treadmill belt was continuously moving except for when the subject was raised from the belt for 5-min of rest, and it was continually maintained at only one speed per subject.

### Stimulation Procedures

#### Tail pinch stimulation

To administer tail pinch, a small amount of gentle pressure was applied about 1 cm from the base of the tail using miniature forceps (Swann et al., 2017). The pressure was very brief, just until reaction to the stimulation was observed, and there was no apparent damage to the tail.

#### Quipazine stimulation

An intraperitoneal injection of quipazine maleate (3.0 mg/kg, 50 microliters, 30-gauge needle) was used to induce alternating stepping. The drug came from Sigma-Aldrich (St. Louis, MO). Quipazine is a 5-HT_2_ receptor agonist and is often used at this dose to induce alternating stepping in neonatal rats (Strain & Brumley, 2014).

### Behavior analysis

All sessions were recorded onto DVD using a camera located in the sagittal plane on the left side of the subject. During DVD playback, stepping and non-stepping movements were scored using the software program JWatcher (Version 1.0) that records the category of behavior and the time of entry (+/- 0.01s). Two behavioral scorers had an intra- and inter-reliability rate of > 90%. Each subject required two scoring passes, in order to score the stepping behaviors of the forelimbs and hindlimbs separately.

#### Step frequency

Alternating stepping was defined as bilateral consecutive alternation of flexion and extension in homologous limb pairs (Brumley et al., 2012). All other movements were scored as non-stepping movements. Only steps in contact with the treadmill were scored. Stepping was analyzed in 30-sec time bins following tail pinch, and in 5-min bins following quipazine administration, due to differences in reaction to the different forms of stimulation.

### Kinematic analyses

For limb kinematic analyses, steps were selected for each part of the test session. As mentioned previously, the tail pinch test session was divided into 30-sec time bins, whereas the quipazine test was divided into 5-min time bins. From each time bin, the first 3-5 consecutive alternating steps were selected for limb kinematic analysis.

#### Step cycle

Swing phase started when the paw of the limb lifted from the treadmill belt and ended just before contact with the belt, after swinging forward. Stance phase started when the paw of the limb came into contact with the treadmill belt and ended just before the paw lifted from the belt to start the swing phase. Total step cycle duration was calculated by adding swing phase and stance phase durations together. Swing, stance, and total step cycle durations were analyzed among groups.

#### Step area

Step area was calculated by multiplying step length and step height (Cha et al., 2007). Step length was measured as the distance of the paw when it was extended the farthest forward and the farthest backward. Step height was measured as the distance of the paw when it reached its highest point and lowest point when in contact with the treadmill belt. Measurements were made by pausing selected frames and marking the points of the limbs on a computer screen. The points were then measured using a ruler, and calibrated using the small ruler attached to the horizontal bar within each video recording.

### Statistical Analysis

Forelimb and hindlimb activity were analyzed separately, using the software program StatPlus v6.1.60. Stepping behaviors and kinematic parameters were grouped into 30-sec time bins for tail pinch stimulation sessions and 5-min time bins for quipazine stimulation sessions. Time-dependent changes in speed-induced stepping and coordination were examined using a repeated measures ANOVA. Independent variables were belt speed and time (the repeated measure). Dependent variables were alternating step frequency, step cycle duration, swing cycle duration, stance cycle duration, and step area. Tukey’s post-hoc comparisons of means and t-tests were performed following findings of significant main effects and/or interactions, respectively. A 5% significance level was used.

## Results

### Effect of Treadmill Speed on Tail Pinch-Induced Stepping: Forelimbs

#### FL (Forelimb) Step Frequency

A two-way repeated measures ANOVA (4 speed groups x 2 30-s time bins) showed a main effect of speed (*F*(3, 28) = 4.80, *p* = .008) and a main effect of time (*F*(1, 28) = 30.38, *p* < .001) on the frequency of alternating forelimb steps following tail pinch. For all speed groups, significantly more alternating forelimb steps occurred within the first 30-s bin compared to the second 30-s bin. Additionally, the medium treadmill speed group showed significantly more forelimb steps compared to the control group (Figure 2A).

**Figure 2.**
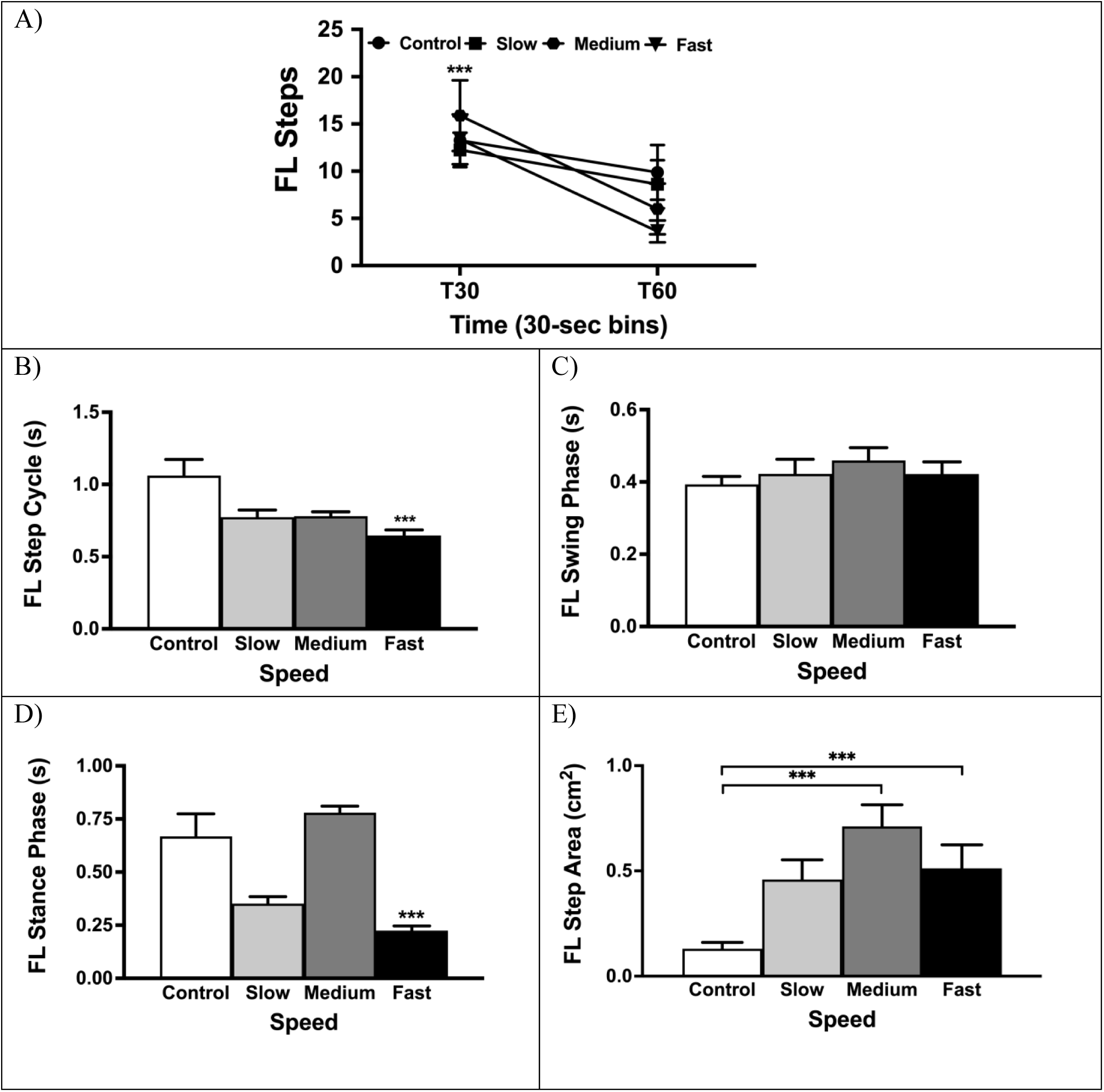
Tail-pinch-induced treadmill stepping in the forelimbs. A). *Frequency of alternating stepping at the first time-bin (T30) and second time-bin (T60).* Subjects showed significantly more forelimb alternating steps in the first time-bin compared to the second time-bin. Subjects showed more alternating steps on the medium belt speed compared to the control belt speed. B). *Forelimb step cycle duration.* Subjects showed significantly shorter step cycle durations on the fast belt speed compared to all other speed groups. C). *Forelimb swing phase duration.* There were no significant differences across speed groups. D). *Forelimb stance phase duration.* Subjects shoed significantly shorter stance phase durations on the fast belt speed compared to all other belt speeds. E). *Forelimb step area*. Subjects on the medium and fast treadmill belt speeds showed significantly larger step areas compared to the control group. Note: *** denotes p < .001.

#### FL Step Cycle

Because very few alternating steps occurred in the second time bin, we analyzed limb kinematics only during the first time bin due to having a sufficient sample size. A one-way ANOVA showed a main effect of speed (*F*(3, 28) = 7.11, *p* < .001) for forelimb step cycle duration. The fast treadmill speed group had significantly shorter step cycle durations compared to the slow, medium, and control speed groups (Figure 2B). For *swing* duration, there were no significant effects (Figure 2C). For *stance* duration, there was a main effect of speed (*F*(3, 28) = 11.43, *p* < .001). The fast group showed significantly shorter stance durations compared to all other groups (Figure 2D).

#### FL Step Area

For forelimb step area, a one-way ANOVA showed a main effect of speed (*F*(3, 28) = 7.10, *p* < .001). The medium and fast groups showed significantly greater forelimb step areas compared to the control group (Figure 2E).

### Effect of Treadmill Speed on Tail Pinch-Induced Stepping: Hindlimbs

#### HL (Hindlimb) Step Frequency

A two-way repeated measures ANOVA showed a main effect of time (*F*(3, 28) = 37.54, *p* < .001) on the frequency of alternating hindlimb steps following tail pinch stimulation. During the first 30-s bin, significantly more alternating hindlimb steps occurred compared to the second 30-s time bin (Figure 3A). There was no effect of treadmill belt speed on hindlimb step frequency following tail pinch.

**Figure 3.**
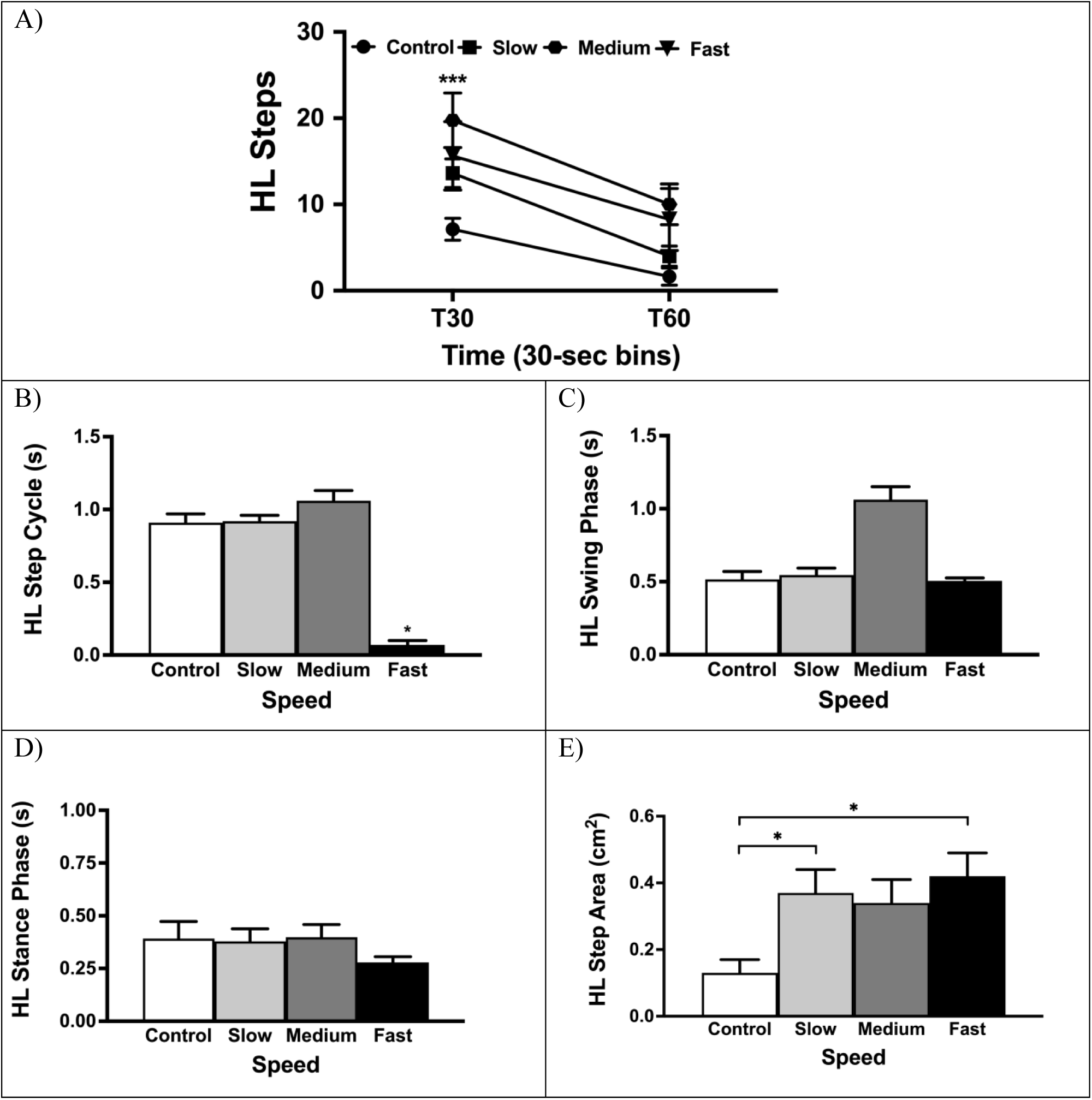
Tail-pinch-induced treadmill stepping in the hindlimbs. A). *Frequency of alternating stepping at the first time-bin (T30) and second time-bin (T60).* Subjects showed significantly more hindlimb alternating steps in the first time-bin compared to the second time-bin. B). *Hindlimb step cycle duration*. Subjects showed significantly shorter step cycle durations on the fast belt speed compared to the slow belt speed group. C). *Hindlimb swing phase duration.* There were no significant differences across speed groups. D). *Hindlimb stance phase duration.* There were no significant differences across speed groups. E). *Hindlimb step area*. Subjects on the slow and fast treadmill belt speeds showed significantly larger step areas compared to the control group. Note: *** denotes p < .001; * denotes p < .05.

#### HL Step Cycle

For hindlimb step cycle duration, a one-way ANOVA showed a main effect of speed (*F*(3, 24) = 4.27, *p* = .015). Subjects in the slow treadmill speed group had significantly longer hindlimb step cycle durations compared to those in the fast group (Figure 3B). For *swing* and *stance* cycle durations, there were no significant effects (Figures 3C & 3D).

#### HL Step Area

A one-way ANOVA showed a main effect of speed (*F*(3, 24) = 3.97, *p* < .02) for hindlimb step area. Subjects in the slow and fast speed groups showed significantly greater step areas compared to those the control group (Figure 3E).

### Summary of Adaptations in Tail Pinch-Induced Stepping

Following tail-pinch, subjects showed significantly more forelimb steps on the medium belt speed compared to the control belt speed. Hindlimb step frequency was not affected by belt speed, but there were significantly more hindlimbs steps in the first 30-s bin compared to the second 30-s bin. When looking at step cycle duration in the forelimbs following tail-pinch, it was found that differences in duration (shorter durations on faster belt speeds) were driven by changes in the stance phase duration. For the hindlimbs, step cycle duration was longest on the slow belt speed compared to the fast speed, but there were no differences in duration of the swing and stance phases following tail-pinch. Step area in the forelimbs was largest in the medium and fast speeds, while the step area for the hindlimbs was largest in the slow and fast speeds.

### Effect of Treadmill Speed on Quipazine-Induced Stepping: Forelimbs

#### FL Step Frequency

A two-way repeated measures ANOVA (4 speed groups x 6 5-min time bins) showed a main effect of speed (*F*(3, 28) = 12.85, *p* < .001) and a main effect of time (*F*(5, 140) = 13.95, *p* < .001) on the frequency of alternating forelimb steps following treatment with quipazine. The control group showed significantly fewer forelimb steps compared to all other groups and the slow treadmill speed group showed significantly fewer steps compared to the fast group. All groups showed significantly fewer forelimb steps in the first 5-min time bin compared to the 15, 20, and 25-min time bins. This can be seen in Figure 4A.

**Figure 4.**
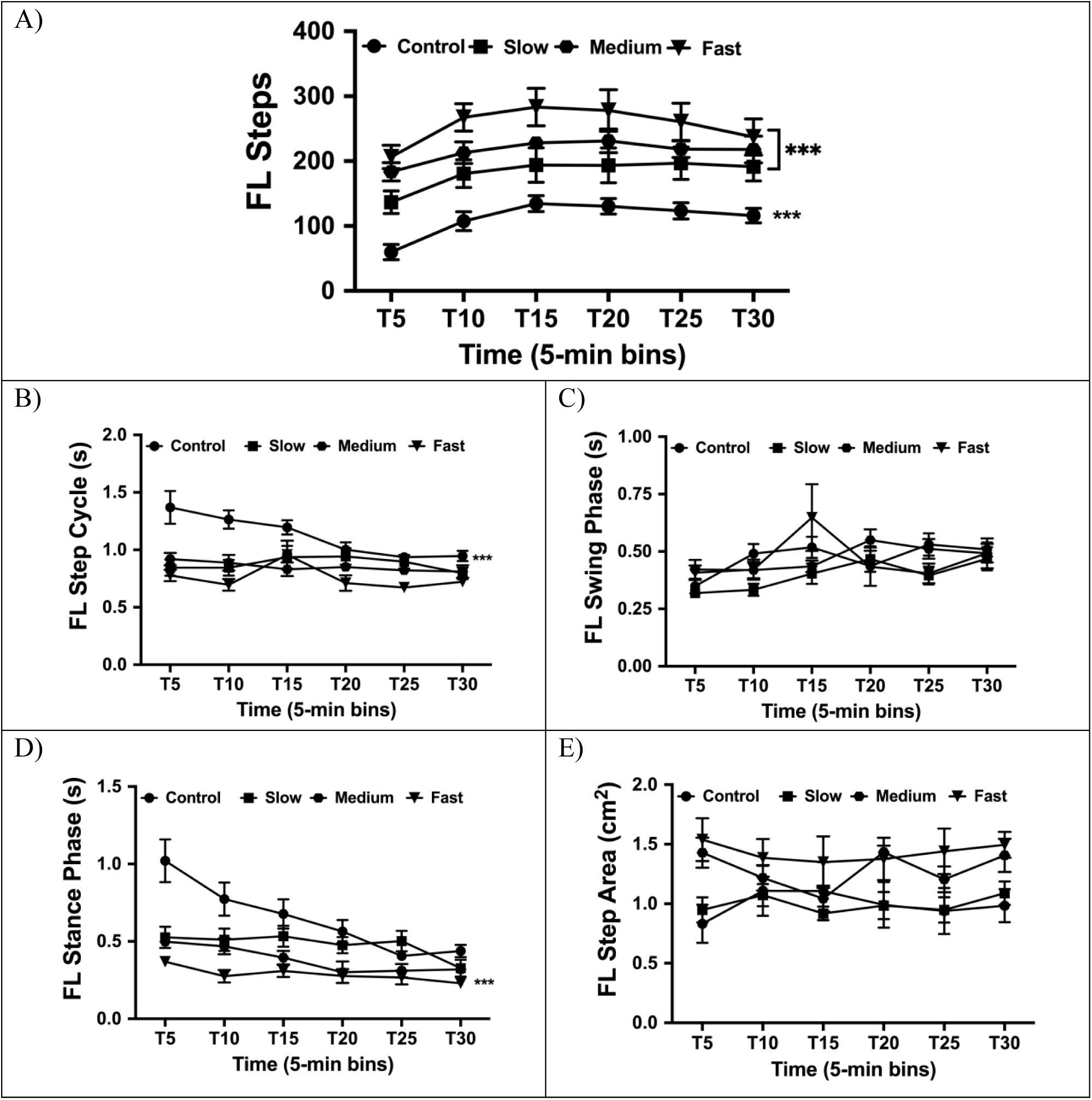
Quipazine-induced treadmill stepping in the forelimbs. A). *Frequency of alternating stepping.* Subjects on the control belt speed show significantly fewer steps compared to all other speed groups. All subjects showed significantly fewer steps in the first time 5-min bin compared to the 15-, 20-, and 25-min bins. B). *Forelimb step cycle duration.* Subjects on the fast belt speed had significantly shorter step cycle durations compared to the control speed at every time bin except T15. C). *Forelimb swing phase duration.* There were no significant differences for speed or time. D). *Forelimb stance phase duration*. Subjects on the fast belt speed had significantly shorter stance phase durations compared to the control group at every time bin except T25. E). *Forelimb step area*. There were no significant differences for speed or time. Note: *** denotes p < .001.

#### FL Step Cycle

For forelimb step cycle duration, a two-way repeated measures ANOVA showed an interaction of speed and time, (*F*(15, 140) = 2.28, *p* = .006), a main effect of speed (*F*(3, 28) = 23.31, *p* < .001), and a main effect of time (*F*(5, 140) = 4.57, *p* < .001). Subjects in the fast treadmill speed group had significantly shorter forelimb step cycle durations compared to subjects in the control group in every time bin except T15 (Figure 4B). For *swing* cycle duration there was a main effect of time (*F*(5, 140) = 3.73, *p* = .003). A one-way ANOVA was used to follow-up the time effect, but it revealed no significant differences between groups. As can be seen in Figure 4C, there was a slight increase in swing duration over time. For *stance* cycle duration there was an interaction of the two factors (*F*(15, 140) = 2.80, *p* < .001), a main effect of speed (*F*(3, 28) = 20.78, *p* < .001), and a main effect of time (*F*(5, 140) = 12.17, *p* < .001). Subjects in the fast treadmill belt speed group had significantly shorter stance durations compared to subjects in the control group in all time bins except T25 (Figure 4D).

#### FL Step Area

For forelimb step area, a two-way repeated measures ANOVA revealed a main effect of speed (*F*(3, 28) = 3.55, *p* = .027). A one-way ANOVA was used to follow-up the main effect of speed, but revealed no significant differences between groups Figure 4E.

### Effect of Treadmill Speed on Quipazine-Induced Stepping: Hindlimbs

#### HL Step Frequency

For hindlimb step frequency, a two-way repeated measures ANOVA showed a main effect of speed (*F*(3, 28) = 12.87, *p* < .001) and a main effect of time (*F*(5, 140) = 24.74, *p* < .001) following treatment with quipazine. Subjects on the control belt speed showed significantly fewer steps compared to all other speed groups. Additionally, subjects on the slow belt speed showed significantly fewer steps compared to the medium and fast belt speed groups. As can be seen in Figure 5A, all groups showed significantly fewer hindlimb steps in the first two 5-min times bins compared to all following time bins.

**Figure 5.**
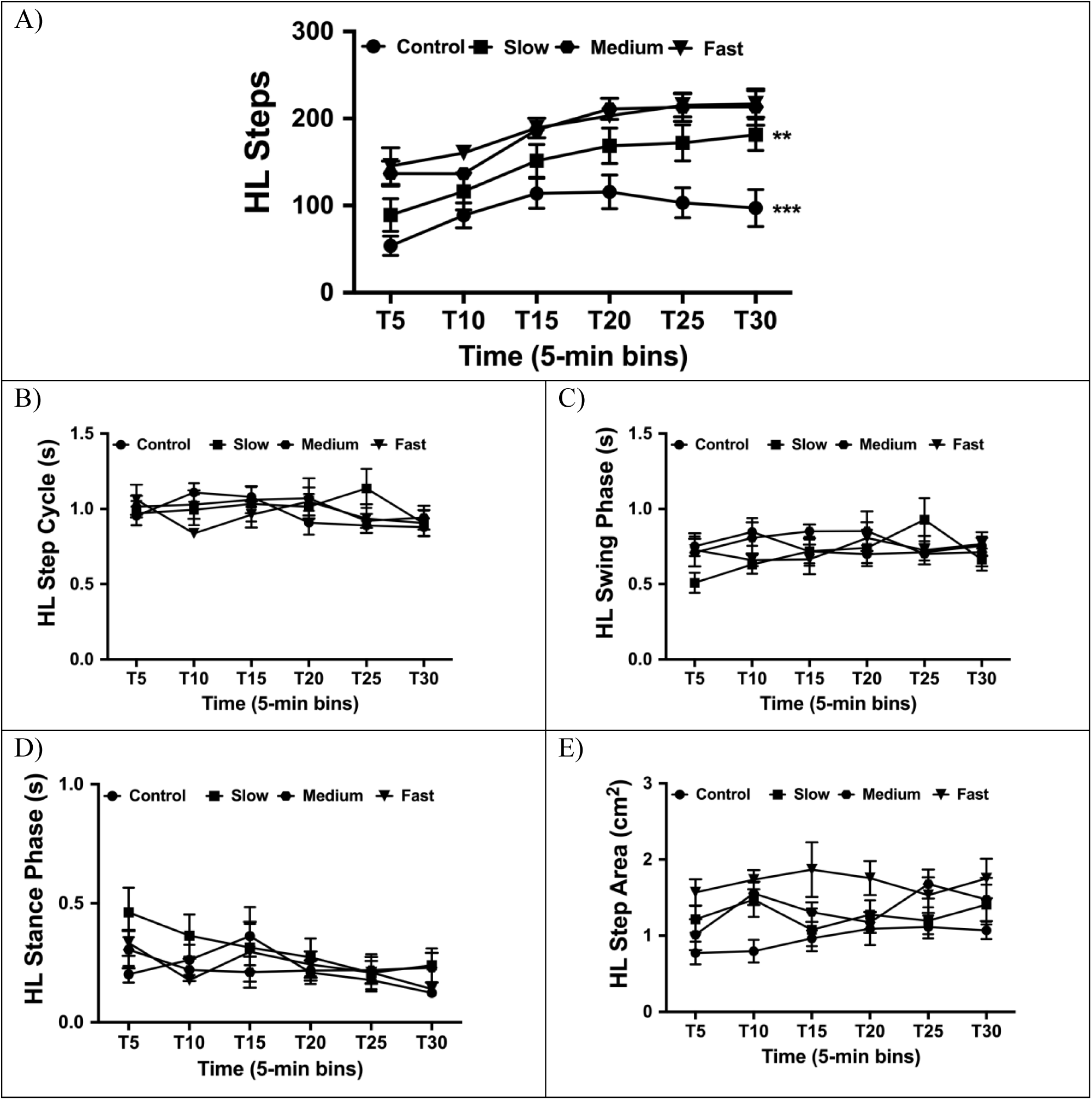
Quipazine-induced treadmill stepping in the hindlimbs. A). *Frequency of alternating stepping.* Subjects on the control belt speed showed significantly fewer steps compared to all other speed groups, and subjects on the slow belt speed showed significantly fewer steps compared to the medium and fast belt speed groups. B). *Hindlimb step cycle duration*. There were no significant differences for speed or time. C). *Hindlimb swing phase duration.* There were no significant differences for speed or time. D). *Hindlimb stance phase duration.* There were no significant differences for speed or time. E). *Hindlimb step area*. There were no significant differences for speed or time. Note: *** denotes p < .001; ** denotes p < .01.

#### HL Step Cycle

For hindlimb step cycle duration and *swing* cycle duration, two-way repeated measures ANOVAs showed no effects of speed or time (Figures 5B & 5C). However, there was a main effect of time (*F*(5, 140) = 2.64, *p* = .026) on hindlimb *stance* cycle duration. A one-way ANOVA was used to follow-up the main effect of time, but showed no significant differences among groups. As can be seen in Figure 5D, there was a slight decrease in stance cycle duration over time.

#### HL Step Area

For hindlimb step area, a two-way repeated measures ANOVA showed only a main effect of speed (*F*(3, 28) = 4.38, *p* = .004). Subjects in the fast treadmill belt speed group displayed significantly larger step areas compared to the control group, as shown in Figure 5E.

### Summary of Adaptations in Quipazine-Induced Stepping

For quipazine-induced stepping, the forelimbs showed the fewest steps in the control belt speed condition compared to all other speed groups. Additionally, there were significantly fewer steps in the first 5-min compared to later time bins. Similarly, the hindlimbs showed fewer steps in the first two time bins compared to later time bins. These time effects are consistent with previous work using quipazine in neonatal rats (Swann et al., 2016). For step cycle, the forelimbs showed the shortest step cycle duration on the fast belt speed compared to the control condition which was driven by changes in durations of the stance phase instead of the swing phase. However, for the hindlimbs, overall step cycle, swing phase, and stance phase durations were not significantly different across belt speeds or time bins.

## Discussion

This is the first study to systematically examine locomotor stepping in newborn rats on a treadmill. Our findings indicate that one-day-old rats are able to step on a treadmill and adapt their step quality in response to a treadmill belt following two forms of stimulation: mechanical (tail-pinch) and pharmacological (quipazine administration).

### Tail-Pinch versus Quipazine-Induced Stepping

Although both paradigms induced alternating stepping as expected, quipazine-induced stepping was much more robust and long-lasting compared to the tail-pinch. Serotonin and its receptor agonists are known to reliably produce locomotion and fictive locomotion (neural rhythm of locomotion) in the newborn rat (Sławińska, Miazga, & Jordan, 2014; McEwen, Van Hartesveldt, & Stehouwer, 1997). Given that quipazine was administered intraperitoneally, the effects of the drug were more systemic, activating neural circuitry throughout the spinal cord and brain, compared to more local activation of spinal circuits with the tail-pinch. The stimulation from the tail-pinch activates sacrocaudal afferents in the spinal cord, which can produce fictive locomotion, but at a lower frequency and for shorter durations compared to the thoracolumbar circuitry (Lev-Tov, Delvolve, & and Kremer, 2000). The whole-body effect of quipazine may be why stepping was much more robust and longer-lasting compared to the induction of stepping from a brief tail-pinch.

While both paradigms induced alternating stepping, these two methods may be acting on different neuronal populations and spinal pathways. Previous research has shown evidence of pathways connecting lumbar central pattern generators (CPGs) and sacrocaudal afferents in neonatal rats, and these sacral networks can be used to activate the hindlimb locomotor CPG (Strauss & Lev-Tov, 2003; Cherniak et al., 2014). Given that a tail-pinch is a localized stimulus, it induces hindlimb stepping through the activation of sacrocaudal afferents, as opposed to a systemic effect of IP pharmacological stimulation (Spear & Ristine, 1981; Swann et al., 2016). IP quipazine administration could induce stepping at any level of the nervous system (i.e., afferents, interneurons, motoneurons), while a tail-pinch is much more local, yet specific, stimuli for induction of stepping. This may also explain why quipazine has a longer lasting effect on stepping compared to the tail-pinch, even though both produce robust responses across time (quipazine maximally activating stepping later in the test session compared to a tail-pinch maximally activating stepping at the beginning of the test session).

### Differences in Forelimb and Hindlimb Stepping Adaptations

Although neonatal rats can show patterns of alternating stepping across many paradigms *in utero* and early in development, there were differences in step quantity and quality in the forelimbs and hindlimbs. Previous research has shown that weight-bearing locomotion can occur early in the postnatal period, but it becomes much more common around P10 in rats (Altman & Sudarshan, 1975; Swann & Brumley, 2019). During the early postnatal period, rat pups engage in forelimb-heavy movements, such as crawling without hindlimb engagement and facial wiping (Altman & Sudarshan, 1975; Eilam & Smotherman, 1998). While the necessary spinal circuitry is in place by birth for producing hindlimb stepping in rat pups, experience with weight-bearing in the hindlimbs may be critical for more mature stepping behavior on a moving treadmill belt (Cazalets et al., 1995). Further, the nervous system and behavior engage reciprocally – that is, neural circuits support behavior and behavioral experience strengthens these pathways (Inglis, Zuckerman, & Kalb, 2000; Tahayori & Koceja, 2012). It is possible that mechanisms of activity-dependent plasticity may drive the development and refinement of hindlimb stepping circuits, thus accounting for the discrepancy in forelimb and hindlimb stepping differences (i.e., forelimbs showing significantly faster stepping on faster speeds, while speed differences in hindlimb stepping were not as expansive) in the present study. This explanation is further supported by past research showing that hindlimb loading (i.e., changes in the amount of body-weight supported during treadmill stepping) affects paw placement, weight-bearing, and movement scores in adult rats spinalized as neonates (Timoszyk, et al., 2005).

The nervous system, particularly the spinal cord, is still developing after birth which may explain why there are differences in forelimb and hindlimb stepping on a moving treadmill belt in our subjects. Both the cervical and lumbar spinal cord is implicated in alternating stepping in the forelimbs and hindlimbs, respectively. The cervical spinal tracts and projections develop earlier in the postnatal period while projections to the lumbar cord that create both ascending and descending tracts are not fully developed until the first and second postnatal weeks (Lakke, 1957). Given that these two sections of the spinal cord that innervate the forelimbs and hindlimbs are considered fully developed at different time periods, it is possible that the necessary circuitry to support hindlimb alternating stepping on a moving treadmill belt is not as mature (or capable of supporting more mature patterns of stepping) as the circuitry in the forelimbs.

### Developmental Differences in Treadmill Stepping

Compared to previous research with adult rodents and treadmill paradigms, our experiment showed both similarities and differences in stepping between early developmental periods and adulthood. As treadmill belt speed increases, intact, adult cats show more steps (faster steps on faster speeds), similar to our results in the forelimbs of neonatal rats (Miller & Van Der Meché, 1975; Frigon et al., 2014). Further, adult mice show shorter step cycles on faster belt speeds (driven by duration changes in the stance phase), which we replicated in our forelimb data of P1 rats (LeBlond et al., 2003). These similarities suggest that early in development, the necessary neural circuitry for forelimb stepping in rats is in place and can be activated via mechanical and pharmacological stimulation. Although quadrupedal locomotion is not common until a couple weeks later, our experiment showed that activating locomotor circuitry in neonatal rats produces alternating stepping in the forelimbs similar to treadmill stepping observed in adult, intact cats (Frigon et al., 2014).

However, there are also some notable differences between alternating treadmill stepping in developing and adult rodents. In our experiment, postural support was necessary for our subjects since it is not established until the second week of development (Brumley, Kauer, & Swann, 2015; Theodossiou et al., 2019). Adults do not need apparatuses for postural support unless the animal is injured (such as spinal cord injury), or postural changes are often being manipulated as an experimental variable (Timoszyk, et al., 2005; de Leon, et al., 2011). Additionally, previous research shows that adult rodents have similarities in stepping of the forelimbs and hindlimbs, while developing rats show differences in step quantity and quality between the forelimbs and hindlimbs (Brumley, Kauer, & Swann, 2015). This may also be due to the circuitry in place at the time of testing (i.e., the adult nervous system is fully mature while the developing rat is still going through many neural changes; Lakke, 1957).

Understanding treadmill stepping in intact, neonatal animals allows researchers to investigate how sensory input can induce and alter motor output. In particular, we examined how both mechanical and pharmacological stimulation induce and modulate stepping during early development. Additionally, past literature has shown the efficacy of using a tail-pinch and quipazine to evoke stepping in neonatal rats across other paradigms (i.e., air-stepping). Our results build on these studies by showing that a tail-pinch or quipazine can be reliably used to induce stepping in neonatal rats in the presence of sensory input (a moving treadmill belt).

Although neonatal rats are a useful model for examining neurobehavioral and developmental plasticity, the equipment necessary for accurate testing is highly limited. Almost all rodent treadmills marketed for biomedical researchers are for use in adult animals, and these treadmills are not developmentally appropriate for use in neonates (i.e., size of treadmill and belt speed). The necessity of tools and equipment that are appropriate for certain developmental periods cannot be understated, and lack of such equipment may slow research progress. Moreover, this equipment is often costly, creating another barrier for researchers. Not only does the present study provide meaningful data about alternating treadmill stepping in neonatal rats, but it also highlights the utility of a developmentally-appropriate treadmill for experimental paradigms. The need for such equipment spurred the design of the treadmill in the present study, as well as the creation of Rat Track (a low-cost, open-source treadmill protocol for use in biomedical models) through collaborative efforts in our lab (Williams et al., 2020).

### Limitations and Future Directions

The current experiment is not without limitations. While the hindlimbs and forelimbs showed similar patterns of effects in each paradigm, the forelimbs showed much more mature locomotion patterns (larger, significant differences between speed groups) compared to the hindlimbs (smaller, nonsignificant differences between speed groups). Although this is the first study to our knowledge to examine treadmill stepping in neonatal rats, past research with adult animals has shown similar trends in step quantity and kinematics between forelimbs and hindlimbs, suggesting that immature, neonatal rat pups show stepping behavior and real-time adaptations when postural support is in place (Frigon et al., 2014; LeBlond et al., 2003). Given that experience with weight-bearing movements, which has also been demonstrated experimentally, may influence more mature patterns of stepping on a moving treadmill belt, testing subjects at later ages, particularly after quadrupedal locomotion is established, may show even more mature patterns of stepping without the need for mechanical or pharmacological induction (Timoszyk et al., 2005).

## Conclusions

This is the first study to examine one-day-old rats stepping on a moving treadmill belt and their ability to adapt stepping in real-time. Overall, we showed that both mechanical (tail-pinch) and pharmacological (quipazine) stimulation is reliable at inducing alternating stepping on a moving treadmill belt in the forelimbs and hindlimbs of neonatal rats. While both stimuli produced effects, we found that quipazine-induced stepping produced prolonged stepping compared to tail-pinch-induced stepping. Further, we showed that P1 rats are capable of modulating both step quantity (frequency) and step quality (kinematics) in response to changes in a moving treadmill belt. Interestingly, the real-time adaptations mimicked stepping adaptations seen in adult rats on moving treadmill belts, suggesting that immature, newborn rats have the necessary circuitry to support step adaptations following stimulation. Examining behavior during developmental periods illuminates the behavioral potential of the developing nervous system and shifts our understanding of behavioral ability in altricial species.

